# Kinetics of Atg2-mediated lipid transfer from the ER can account for phagophore expansion

**DOI:** 10.1101/2020.05.12.090977

**Authors:** Sören von Bülow, Gerhard Hummer

## Abstract

The protein Atg2 has been proposed to form a membrane tether that mediates lipid transfer from the ER to the phagophore in autophagy. However, recent kinetic measurements on the human homolog ATG2A indicated a transport rate of only about one lipid per minute, which would be far too slow to deliver the millions of lipids required to form a phagophore on a physiological time scale. Here, we revisit the analysis of the fluorescence quenching experiments. We develop a detailed kinetic model of the lipid transfer between two membranes bridged by a tether that forms a conduit for lipids. The model provides an excellent fit to the fluorescence experiments, with a lipid transfer rate of about 100 per second and protein. At this rate, Atg2-mediated transfer can supply a significant fraction of the lipids required in autophagosome biogenesis. Our kinetic model is generally applicable to lipid-transfer experiments, in particular to proteins forming organelle contact sites in cells.

## INTRODUCTION

Atg2 has been proposed to function as a conduit for the transfer of phospholipids from the ER to the phagophore in autophagy (Chowdhury et al. 2018, Osawa et al. 2019, Valverde et al. 2019). Osawa et al. (2019) recently reported the crystal structure of the N-terminal region of *Schizosaccharomyces pombe* Atg2. A hydrophobic groove along the long axis of Atg2 could accommodate phospholipid acyl chains, in resemblance to other lipid-transfer proteins. By mixing lipid vesicles with and without fluorescently labeled lipids, Osawa et al. (2019) also demonstrated transfer of lipids between vesicles in an Atg2-dependent manner. The crystal structure of Atg2 (Osawa et al. 2019) and independent lipid transfer measurements by several groups (Maeda et al. 2019, Osawa et al. 2019, Valverde et al. 2019) provide strong evidence that Atg2 tethers membranes and acts as a lipid-transfer protein to supply phospholipids during autophagosome formation.

However, recent kinetic measurements put into question that the bulk of lipids in phagophores passes through Atg2. From careful fluorescence measurements, Maeda et al. (2019) deduced a transfer rate of only 0.017 labeled lipids per second and ATG2A, which is the human homolog of yeast Atg2. Phagophores of typical size contain on the order of 10^6^ lipids. At the reported transfer rate of 0.017 s^−1^ and with Atg2 as the sole conduit, it would take years to form a phagophore. Clearly, at this rate Atg2 cannot account for phagophore expansion, which occurs on a time scale of minutes in living cells. Even a 50-fold increase, proposed as a possible correction for the low abundance of labeled lipids, does not resolve this issue. Therefore, Maeda et al. (2019) concluded that other factors must be at play.

Here, we revisit the analysis of the kinetic measurements by Maeda et al. (2019). We show that the peculiar aspects of single-file transport (Berezhkovskii and Hummer 2002, Hummer et al. 2001, Kalra et al. 2003) in the hydrophobic groove acting as lipid conduit of Atg2 require a reinterpretation of the kinetic measurements. In single-file transport, the transferred molecules (here: lipids) are arranged in a one-dimensional chain. This chain moves back and forth in a stochastic manner, and thereby releases molecules into the reservoir at one end and picks up molecules from the reservoir at the other end. Through a number of such back-and-forth hopping events, molecules are transferred from one reservoir to the other. See Figure 1 for a schematic representation of a minimal lipid transfer model. Two central aspects of single-file transport are that all transferred molecules pass through a single bottleneck and that the elementary event (here the back-and-forth hopping motion of the chain of lipids) occurs stochastically with a given rate (or diffusion coefficient). As a consequence, the flux through the bottleneck is limited to about one molecule per characteristic time more or less independent of the abundance of the transported molecules in each reservoir.

**Figure 1.**
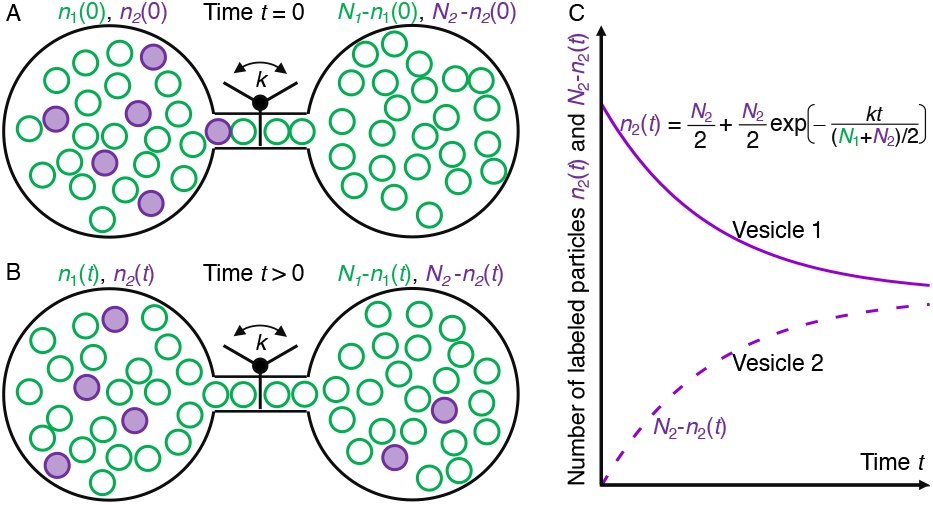
Minimal model of lipid transfer between two equal-size vesicles. (A) At time *t* = 0, vesicle 1 (left) contains *n*_1_(0) unlabeled lipids (green) and *n*_2_(0) = *N*_2_ labeled lipids (magenta). Vesicle 2 contains *n*_1_ − *n*_1_(0) unlabeled lipids and *N*_2_ − *n*_2_(0) = 0 labeled lipids. The two vesicles are connected by a channel that mimics an Atg2 tether. At a rate *k*, single lipids are shuttled from one vesicle to the other in a random direction and irrespective of the presence of a fluorescence label. This single-lipid transfer through the channel is akin to a turnstile rotating stochastically by 120° either clockwise or anti-clockwise at frequency *k*. On the vesicles, we assume that lipid mixing is fast compared to lipid transfer. (B) After time *t*, mostly unlabeled lipids have transferred, but also some labeled lipids. (C) As a result of lipid transfer, the populations of labeled lipids on the two vesicles relax exponentially with a rate *r* = *k/N* to their limiting value *N*_2_/2, where *N* = (*N*_1_ + *N*_2_)/2 is half the total number of lipids.

As a consequence, the apparent rate *r* of the lipid-population relaxation in fluorescence quenching assays (Maeda et al. 2019, Osawa et al. 2019, Valverde et al. 2019) has to be multiplied by the total number *N* of lipids in a leaflet of the Atg2-tethered vesicles to get the rate of lipids passing through Atg2, *k* = *rN*. We present a detailed kinetic model of the transport and solve the corresponding full master equation analytically. Using Förster theory, we also derive expressions for the expected time dependence of the donor fluorescence intensity under the assumption that transported lipids mix rapidly. We then fit these expressions to the fluorescence time traces measured by Maeda et al. (2019). In this way, we estimate a rate of transfer of about 115 lipids per second and ATG2A, about 7,000 times faster than the original estimate. At this rate and with a minimal thermodynamic driving force, ATG2A is capable of transferring a significant fraction of the phospholipids from the ER and/or lipid droplets to the growing phagophore. In fits to fluorescence time traces for yeast Atg2 (Osawa et al. 2019), we obtain an even faster rate of about 750 lipids per second and Atg2. In the cell, Atg2mediated lipid transfer is expected to be modulated by lipid composition, membrane curvature and protein co-factors, in particular WIPI4/Atg18 (Chowdhury et al. 2018, Maeda et al. 2019, Osawa et al. 2020, 2019).

## RESULTS

The Förster resonance energy transfer (FRET) assay used to study protein-mediated lipid transport between vesicles (Connerth et al. 2012, Kawano et al. 2018, Maeda et al. 2019, Osawa et al. 2019, Valverde et al. 2019, Watanabe et al. 2015) is a variant of an assay for vesicle-vesicle fusion (Struck et al. 1981). In the lipid transfer experiments of Maeda et al. (2019), vesicles containing a small fraction of lipids labeled with fluorescence dyes were mixed with unlabeled vesicles. As fluorescence donors and acceptors, Maeda et al. (2019) used 2% dioleoyl phosphatidylethanolamine nitrobenzoxadiazole (NBD-PE) and 2% dioleoyl phosphatidylethanolamine lissamine rhodamine (rhodamine-PE) lipids, respectively, in a background of 46% dioleoyl phosphatidylcholine (DOPC), 25% dioleoyl phosphatidylethanolamine (DOPE) and 25% dioleoyl phosphatidylserine (DOPS). The unlabeled vesicles contained 50% DOPC, 25% DOPE and 25% DOPS. After mixing the lipids in the presence of Atg2 at different concentrations, Maeda et al. (2019) monitored the change in fluorescence as a function of time. From these time traces, they deduced the rate of lipid transfer.

We applied the analytical expression for the fluorescence signal in Atg2-mediated lipid transfer between vesicles developed in the Theory section below to the measurements of Maeda et al. (2019). We focus on the ATG2A concentration-dependent data in their Figure 3C. For simplicity, we assumed that all vesicles have a uniform diameter of 50 nm and thus an outer radius of *R*_1_ = 25 nm. This diameter corresponds to the peak in the vesicle-size distribution Figure 3–Figure Supplement 1 of Maeda et al. (2019). However, the reported size distributions are for vesicles of somewhat different lipid composition from the ones used in the lipid transfer experiments. Below, we will also discuss the effect of varying the vesicle size in the model. To determine the number of lipids in each leaflet, we used an area per l ipid of 0.713 n m^2^, a value obtained by linear interpolation to 25°C of the values reported for DOPC (Pan et al. 2008). We arrive at *N* ≈ 11,000 lipids in total in each of the outer leaflets, and ≈ 7,400 lipids in the inner leaflets for a membrane thickness of *h* = 4.5 nm, corresponding to an inner radius of *R*_2_ = *R*_1_ *− h* = 20.5 nm. A 2% fraction then results in 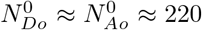 NBD-PE and rhodamine-PE lipids in the outer leaflets of labeled vesicles, and 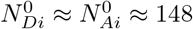 lipids in the inner leaflets.

For the FRET calculations, we used a Förster radius of *R*_0_ = 4.6 nm for the NBD-PE and rhodamine-PE FRET pair (Lantzsch et al. 1994). For the outer and inner vesicle radii *R*_1_ and *R*_2_, and the given Förster radius of the fluorophore pair, the quenching factors are *Q_oo_ ≈* 0.010, *Q_oi_ ≈* 0.004 and *Q_ii_ ≈* 0.015, as calculated from Equation 25.

As a first step, we modeled the background signal in Figure 3C of Maeda et al. (2019), as obtained in the absence of ATG2A. Over the time range of the experiment, we could fit this signal well with a single exponential, *b*(*t*) = 4.28 × [1 − exp(−*r_b_t*)] with *r_b_* = 0.001 88 s^−1^ the rate of increase in the background signal. See data and fit for “0 nM” in Figure 2A.

**Figure 2.**
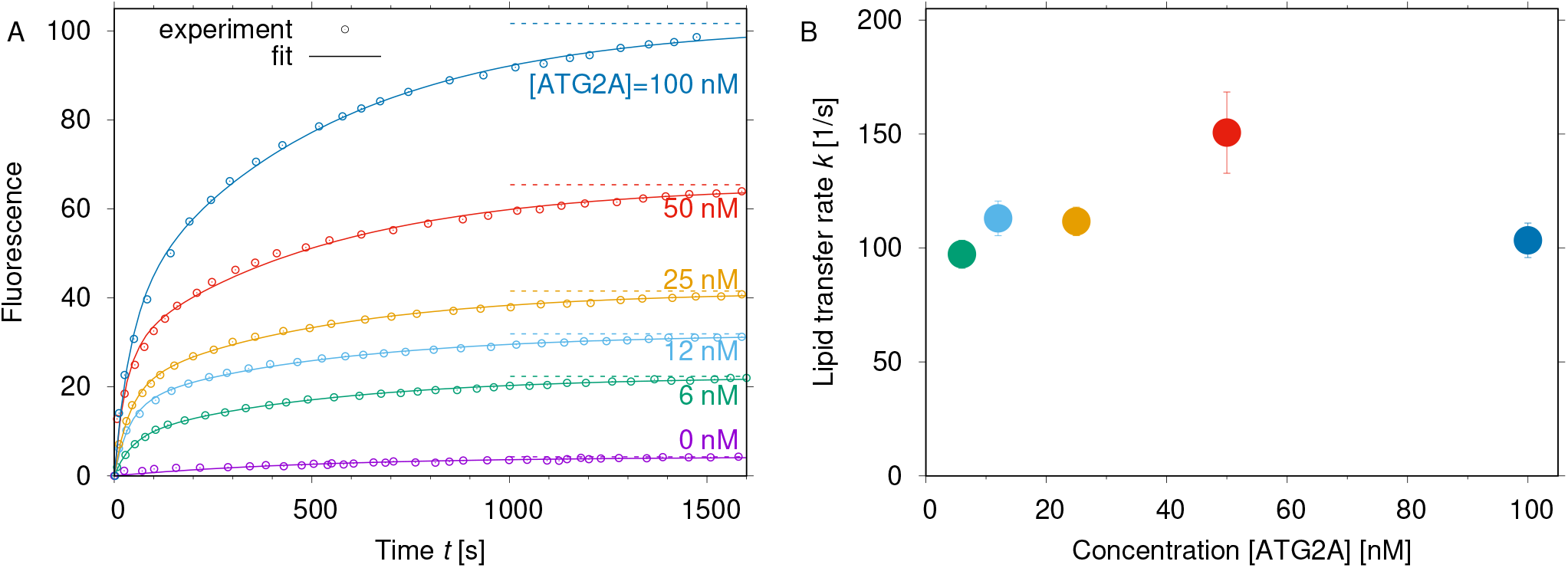
Rate of ATG2A-mediated lipid transfer from fit of kinetic model to FRET lipid-transfer measurements by Maeda et al. (2019). (A) Fit of two-leaflet kinetic model to FRET lipid-transfer measurements. The experimental data (symbols) for ATG2A-mediated lipid transfer were taken from Figure 3C of Maeda et al. (2019). The solid lines are fits of the kinetic model Equation 30, which includes both intraleaflet and interleaflet FRET. The measured signal without ATG2A (reproduced here and labeled 0 nM) was used as background. Fit parameters were the rate *k* of lipid transfer and the amplitudes of signal and background. Horizontal dashed lines indicate the limiting values as time goes to infinity. (B) ATG2A-mediated lipid transfer rates. The lipid-transfer rates *k* obtained from the fits shown in (A) are plotted as a function of ATG2A concentration. Error bars indicate relative fit uncertainties; i.e., they do not account for statistical uncertainties or for uncertainties in the vesicle diameters and other parameters of the model.

**Figure 3.**
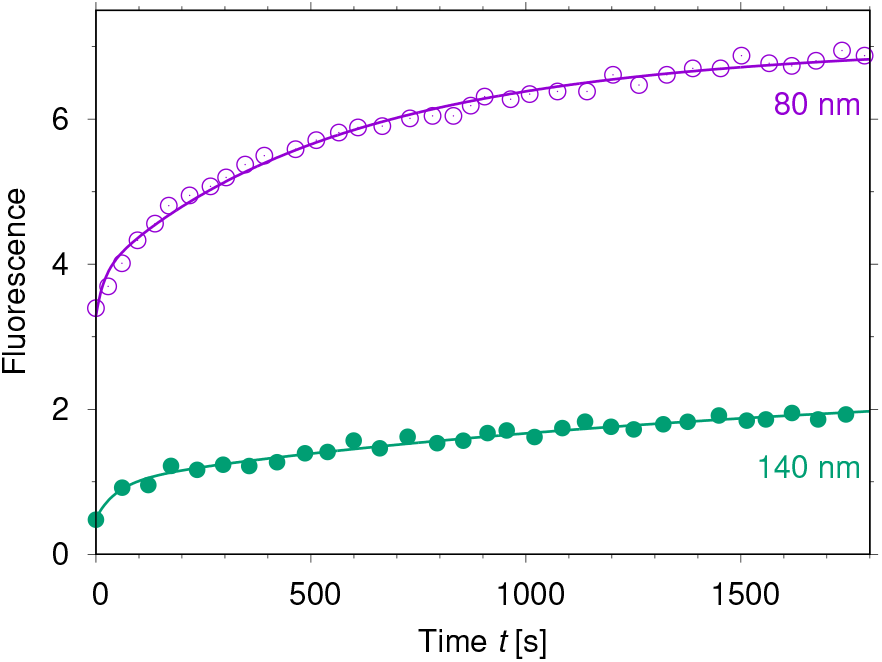
Rate of yeast Atg2-mediated lipid transfer from global fit of kinetic model to FRET lipid-transfer measurements by Osawa et al. (2019). Global fits of the two-leaflet kinetic model with two kinetic populations are shown as lines, and experimental fluorescence traces taken from Figure 3f of Osawa et al. (2019) as symbols. Results are shown for small vesicles (80 nm diameter; magenta) and large vesicles (140 nm diameter; green).

In a second step, we extracted the lipid transfer rate *k* by fitting the measured fluorescence as a function of time for different ATG2A concentrations (Figure 2). We note that the measured fluorescence values reported in Figure 3C of Maeda et al. (2019) were scaled by the value of the fluorescence measured for each condition after detergent-induced dissolution of the lipid vesicles. We reasoned that in the measurements at different concentrations of ATG2A, the background signal will have different relative intensities but the same time dependence. Therefore, we fitted 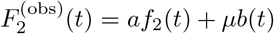 to the fluorescence time traces using the amplitudes *a* and *μ*, and the lipid transfer rate *k* as the fit coefficients, with *f*_2_(*t*) given in Equation 30. Lipid exchange in this model is limited to the outer leaflets, but the calculated fluorescence signal includes FRET also between inner and outer leaflets. The amplitudes *a* and *μ* of signal and background, respectively, are shown in Figure S4. As shown in Figure 2A, we obtained consistently excellent fits to the fluorescence time traces in Figure 3C of Maeda et al. (2019). In particular, the model captures the slightly non-exponential time dependence.

The lipid transfer rate *k* extracted from these fits is independent of the concentration of ATG2A, as shown in Figure 2B, over the entire concentration regime from 6 nM to 100 nM. Treating the five measurements as independent and normal distributed (*p*-value of Shapiro-Wilk test for normality is 0.12), we arrive at a mean value of *k ≈* 115 s^−1^. This rate differs from the apparent rate *r* by a factor *N ≈* 11,000, i.e., by the number of lipids in an outer leaflet. Indeed, the apparent relaxation rate *r* = *k/N* ≈ 0.010 s^−1^ is very close to the value reported by Maeda et al. (2019). To obtain the lipid transfer rate *k*, the apparent rate *r* should be scaled by the number of lipids *N*, not by the reciprocal fraction of labeled lipids.

We also fitted the single-leaflet model *f*_1_(*t*) defined in Equation 24 for *R* = 25 nm (Figure S5G,H). This simplified model ignores interleaflet FRET. With a Förster radius comparable to the membrane thickness, *R*_0_ ≈ *h*, intraleaflet FRET dominates. Accordingly, the fits of the single-leaflet model are indistinguishable from those of the two-leaflet model shown in Figure 2A, and the resulting lipid transfer rates change by less than 1%.

As a word of caution, we want to emphasize that the estimate of *k* ≈ 115 s^−1^ gives us only an order of magnitude of the lipid transfer rate. As discussed below, the sensitivity of the inferred value of *k* to the vesicle radius results in significant uncertainties.

## DISCUSSION

### Plausibility checks

One may wonder how the ATG2A-mediated transfer rate deduced from the same experimental data set could differ so dramatically between, here, 115 per second, and 0.017 per second in (Maeda et al. 2019). So, as a first test, we perform a plausibility check. With 2% donor and 2% acceptor lipids and a diameter of about 50 nm, the labeled vesicles in the fluorescence quenching experiments contain 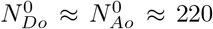 fluorescence donor and acceptor lipids each in the outer leaflet of the labeled vesicles. At the halflife, when the signal has reached 50% of the plateau value, we expect that 2 × (220 − 110)/2 = 110 donor and acceptor lipids have transferred to the outer leaflet of the unlabeled vesicle. Even if this transfer occurred without any lipid back-flow and if only labeled lipids were transferred, one by one, at a frequency of 0.017 s^−1^, the transfer of 110 lipids would take 110/(0.017 s^−1^) ≈ 6500 s. Backflow and transfer of the more abundant unlabeled lipids would increase this time substantially. However, the fluorescence traces in Figure 3C of Maeda et al. (2019) reach 50% of their respective plateau values already after about 200 s (see Figure 2A). On the basis of this simple argument, we conclude that the ATG2A-mediated lipid transport must have occurred with a rate of at least 1 s^−1^. The kinetic model leading to Equation 30 accounts for the abundances of labeled and unlabeled lipids, for the stochastic back-and-forth transport through an Atg2 conduit, and for the dependence of the fluorescence intensity on the density of donor and acceptor lipids in each leaflet.

As a second plausibility check, one may compare Atg2 to bona fide lipid-transfer proteins. Osawa et al. (2019) find that the speed of Atg2-mediated lipid transfer exceeds that of the Mmm1-Mdm12 complex (Kawano et al. 2018) and of VPS13 (Kumar et al. 2018), two protein systems with demonstrated lipid-transfer activity at ER contact sites. Any biologically relevant lipid transfer at an organelle contact site would be expected to be substantially faster than 0.017 s^−1^, i.e., one lipid per minute.

One can also ask whether a rate of *k* ≈ 115 s^−1^ is meaningful at a molecular scale. Following the theory of Berezhkovskii and Hummer (2002) for the kinetics of single-file transfer, as outlined below, and using the reported number of lipids forming the single-file chain, *M ≈* 20 (Valverde et al. 2019), we estimate the elementary hopping rate of the single-file chain as *k*_hop_ = (*M* + 1)*k* ≈ 2400 s^−1^. The characteristic time between hopping events of 1*/k*_hop_ ≈ 410 *μ*s would appear to be reasonable for a complex molecular process. For comparison, the protein TMEM16 accelerates the rate of lipid scrambling between the inner and outer leaflets to greater than 10^4^ lipids per protein and second based on fluorescence bleaching experiments (Malvezzi et al. 2018). Here it is worth noting that in the kinetic analysis of their lipid scrambling experiments, Malvezzi et al. (2018) also scale the apparent rates by the total lipid number to estimate the lipid scrambling rate per protein. Indeed, in TMEM16-mediated scrambling, the lipids have been shown to pass through a bottleneck in a single-file like arrangement (Lee et al. 2018, Siggel et al. 2019).

### Lipid transfer rates are independent of ATG2A concentration

The excellent fit of the theoretical model to the measured curves with a minimal number of adjustable parameters (i.e., the lipid transfer rate *k* and the amplitudes of signal and background) give some reassurance that the kinetic model Equation 30 captures at least the essence of the lipid transport kinetics. Further reassurance comes from the fact that the lipid transfer rate *k* is independent of ATG2A concentration up to 100 nM. By contrast, the rate extracted in the original analysis dropped by more than a factor of three when going from the lowest (6 nM) to the highest ATG2A concentration (100 nM; see Figure 3D in (Maeda et al. 2019)). Such a slowdown with increasing concentration is rather unexpected. If anything, one would expect an increase in the lipid transfer rate because at higher concentrations multiple ATG2A could bridge between labeled and unlabeled vesicles. Indeed, if *m* ATG2A bridges between the vesicles operate in parallel and independent of each other, the lipid transfer rate *k* and the population relaxation rate *r* in the kinetic model based on Equation 1 are accelerated by a factor *m*. The concentration independence of the transfer rate *k* and the net increase in the transfer of labeled lipids together indicate that labeled and unlabeled vesicles are typically bridged by at most one ATG2A protein (i.e, *m* = 0 or 1) and that the bridged population (*m* = 1) increases with concentration.

### Variations in vesicle size

In our fit to the data of Maeda et al. (2019), we assumed that the labeled and unlabeled lipid vesicles have a uniform diameter of 50 nm. However, according to Figure 3–Figure Supplement 1 of Maeda et al. (2019), the vesicle diameters reported for a related experiment with a different lipid composition vary from about 25 nm to 110 nm. Importantly, when the diameter changes but the concentrations of fluorescence donor and acceptor lipids are constant, the product 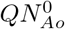 of the quenching factor *Q* and the number 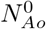 of acceptor lipids stays nearly constant, as follows from the dominant quadratic dependence of *Q* on the ratio *R*_0_*/R* in Equation 20. However, the relaxation rate *r* changes as *r ∝* 1*/ N* ∝ 1*/R*^2^. With 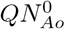 constant and *N* variable, the normalized fit function (and therefore the fit quality) stays essentially unchanged except for a change in time scale. For vesicles with a diameter of 100 nm, the fits to the fluorescence traces in Figure 2A remain unchanged, and the rate *k* of ATG2A-mediated lipid transfer in Figure 2B increases by a factor (100/50)^2^ to ≈ 460 s^−1^ (Figure S5A,B); for vesicles with 25 nm diameter, the rate *k* decreases by a factor (25/50)^2^ to ≈ 30 s^−1^ (Figure S5C,D). Whereas these values are of the same order of magnitude, the precise value of the lipid transfer rate *k* deduced from the fits to the experiments is sensitive to the values of the vesicle diameter.

In a mixture of vesicles of different sizes, larger vesicles with more labeled lipids will contribute more to the signal. We estimated the effect of having a mixture of vesicles by combining the expected signals for vesicles with diameters of 25 nm, 50 nm and 100 nm at relative weights of 25%, 50% and 25%. For simplicity, we assumed that vesicle pairs are tethered by ATG2A at random irrespective of their size. Using the rates *r* for unequal vesicle radii in Equation 15, we averaged the fluorescence signal over the distribution of vesicle sizes. Compared to Figure 2, the fit is slightly improved at short times, and the rate *k* of lipid transfer increases to about 180 lipids per second and ATG2A (Figure S5E,F). We conclude that at least part of the dispersion in the fluorescence signal can be attributed to having a mixture of vesicle sizes.

In this context, one may also want to consider uncertainties in the Förster radius *R*_0_. In fits of the two-leaflet model for a vesicle diameter of 50 nm, a 10% increase/decrease in *R*_0_ results in about a 20% decrease/increase in the rate *k*. Within a reasonable range of uncertainty, changing *R*_0_ does not significantly affect the magnitude of the lipid transfer rate.

### Background signal

We modeled the background fluorescence intensity change according to the fluorescence trace measured in the absence of ATG2A (labeled 0 nM in Figure 3C of Maeda et al. (2019)). Importantly, the apparent rate of increase in the background signal, *r_b_* ≈ 0.001 88 s^−1^ is about ten times slower than the apparent rate *r* = *k/N* ≈ 0.0175 s^−1^ of the increase in the fluorescence s ignal d ue to ATG2A-mediated lipid transfer. Because of this kinetic separation, one can relatively easily distinguish signal and background in the fit. Nevertheless, at relative amplitudes of 20 to 35%, the background signal is appreciable (Figure S4). For a full quantification of the lipid transfer rate, it will therefore be important to characterize the sources of the background signal such as fluorophore bleaching and, if possible, minimize its contribution.

As an alternative, the slow phase in the fluorescence traces is the result of a second population of protein-tethered vesicles with a slower lipid transfer rate. To test this hypothesis, we performed a global fit of a model with two kinetic populations, 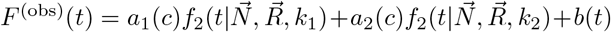 to the FRET lipid-transfer measurements reported in Figure 3C of Maeda et al. (2019). The concentration-dependent amplitudes *a*_1_(*c*) and *a*_2_(*c*) of the two kinetic populations and their concentration-independent lipid-transfer rates *k*_1_ and *k*_2_ were optimized, with *c* = [ATG2A] the concentration of ATG2A. In the global fit, the background signal *b*(*t*) was fixed corresponding to the fluorescence trace without ATG2A (0 nM). The vesicle diameter was set to 50 nm, as in Figure 1, and the two-leaflet model was used.

The results of this global fit are shown in Figure S6. We obtained lipid-transfer rates of *k*_1_ = 142 s^−1^ and *k*_2_ = 10 s^−1^. The amplitudes of the fast and slow phase are comparable in magnitude and increase with ATG2A concentration. The fast rate *k*_1_ agrees well with the average rate *k* ≈ 115 s^−1^ obtained in Figure 1. The slow rate *k*_2_ is about one order of magnitude slower. As pointed out by Osawa et al. (2019), displacement of helix H4 from the hydrophobic groove is likely required for efficient lipid transfer. A sub-population with H4 not fully displaced could account for the slow phase, as could variations in membrane tethering or assembly of ATG2A, or tethered networks of vesicles. Below, we also explore the possibility that lipid flip-flop causes the slow phase. However, the apparent rate *k*_2_*/N* corresponding to the slow transfer rate is close to the rate we estimated for the background, *k*_2_*/N* ≈ 0.001 s^−1^ ≈ *r_b_* = 0.001 88 s^−1^. This close correspondence makes it difficult to separate the slow phase from a background signal of variable amplitude. Nevertheless, a significant fraction of the lipids is transferred at rates exceeding 100 lipids per second irrespective of the model. The evidence for fast lipid transfer is thus strong.

### Apparent kinetics of fluorescence signal depends on vesicle size

In their Figure 3f, Osawa et al. (2019) report lipid-transfer measurements mediated by yeast Atg2 for vesicles of different size. They observed a slower activity for large liposomes compared to small liposomes with diameters of *>* 140 nm and *<* 80 nm, respectively. Whereas the curvature-dependent Atg2 binding affinity to lipid membranes is the dominant factor (Osawa et al. 2019), we hypothesized that at least parts of the observed slowdown in the apparent rate of increase in the fluorescence can be attributed to the fact that the apparent rate *r* scales inversely with the number *N* of lipids per leaflet.

To test this hypothesis, we performed global fits of our theory to the measurements reported in Figure 3f of Osawa et al. (2019) for small and large liposomes and with background already subtracted. Remarkably, a significant fraction of the lipid transfer occurred during the dead time of the experiment. For the 80-nm vesicle, this fraction is nearly 50%. Indeed, the authors point out a time lag in the preparation of reaction mixtures. We account for this lag by a time delay of Δ*t* = 30 s. I.e., our fit function was *F* ^(obs)^(*t* + Δ *t*). For shorter delays Δ*t*, the fast rate *k*_1_ becomes even faster to account for the mostly unresolved rise in intensity. We set the vesicle diameters to 80 nm and 140 nm, respectively, corresponding to the peaks in the reported size distributions.

As shown in Figure 3, a model with two kinetic populations, 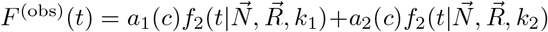, gives very good agreement with the experimental data of Osawa et al. (2019) for Atg2-mediated lipid transfer. For the fast rate of yeast Atg2-mediated lipid transfer, we obtain *k*_1_ ≈ 750 s^−1^. The slow rate is *k*_2_ ≈ 19 s^−1^, with a relative amplitude of the fast phase of 50% and 38% for the 80-nm and 140-nm vesicles, respectively. Irrespective of the cause of the slow phase, the rates *k*_1_ and *k*_2_ obtained for yeast Atg2 and human ATG2A are similar in magnitude. This consistency of the lipid-transfer rates indicates that both proteins operate with a common mechanism.

### Lipid selectivity

We assumed that Atg2 transfers all lipids irrespective of type. For the present experiments, this assumption would appear to be reasonable because labeled and unlabeled lipids have dioleoyl acyl chains, and it is the acyl chains that interact primarily with Atg2 (Osawa et al. 2019). Indeed, (Osawa et al. 2019) concluded that Atg2 exhibits little head-group specificity in its lipid transfer activity. To account for such selectivity, one could make the rate *k* in the rate definitions for 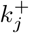 and 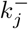 dependent on lipid type, i.e., by using *k_j_* instead of *k*.

### Lipid flip-flop

We also assumed that lipid exchange occurs only between the outer leaflets of the vesicle membranes. For fast lipid flip-flop between the leaflets mediated by an efficient scramblase (Malvezzi et al. 2018), *N* would effectively double. By contrast, if lipid flip-flop between the two leaflets is slow compared to the relaxation rate *r*, we expect the appearance of a second slow phase according to the rate model Equation 33. Therefore, we hypothesized that lipid flip-flop could account for the slow phase.

In Figure S7A and B, we show the results of global fits using a model with lipid flip-flop, Equation 33, to the size-dependent data of Osawa et al. (2019) for Atg2 and to the concentration-dependent data of Maeda et al. (2019) for ATG2A, respectively. In the global fit to the Atg2 data for two vesicle sizes (Osawa et al. 2019), we again used a delay of Δ*t* = 30 s, resulting in a lipid transfer rate of *k* = 336 s^−1^ and a flip-flop rate of *s* = 1.0 × 10^−3^ s^−1^ per lipid. In the global fit to the ATG2A data (Maeda et al. 2019), we fixed the background at *b*(*t*), resulting in *k* ≈ 51 s^−1^ and *s* ≈ 1.1 × 10^−3^ s^−1^, respectively. The fits are considerably worse than those in Figure 1 and Figure 3. Moreover, the flip-flop rates *s* are about 30 times faster than a rate of 3 × 10^−5^ s^−1^ measured for NBD-PE in PC membranes (Armstrong et al. 2003). Overall, factors such as imperfect tethering or assembly, including incomplete H4 helix displacement, appear more likely to cause the slow phase than lipid flip-flop.

### Biased transfer of lipids from the ER to the phagophore

The question driving this research and the preceding studies concerns the origin of the lipids forming the phagophore. Atg2 has been proposed as a conduit for lipid transfer from the ER. As shown by Kalra et al. (2003) for single-file transfer through a nanopore, even a slight thermodynamic bias results in a rapid net transfer. Let Δ*μ* = *μ*_phagophore_ − *μ*_ER_ be the difference in chemical potential (Gibbs free energy) between a lipid on the phagophore and in the ER, and *β* = 1/(*k*_B_*T*) the inverse temperature *T* with *k*_B_ Boltzmann’s constant. The ratio of probabilities for a transfer event to be from the ER to the phagophore (*p*) and from the phagophore to the ER (*q*) is then *p/q* = exp(−*β*Δ*μ*). The expected net number of lipids transferred from the ER to the phagophore during time *t* is Δ*n*(*t*) = *kt*(*p − q*) = −*kt* tanh(*β*Δ*μ/*2) (Kalra et al. 2003). The variance grows linearly with time, var(Δ*n*(*t*)) = *kt*. For a bias as small as Δ*μ* = 1 *k*_B_*T* = 2.5 kJ/mol, a single bridging ATG2A molecule would in ten minutes transfer about Δ*n*(*t* = 10 min) ≈ 32,000 lipids from the ER to the expanding phagophore at a rate of *k* ≈ 115 s^−1^, as determined from the fits to the experiments by Maeda et al. (2019). In the biological setting of phagophore expansion, it will be interesting to study the energetic drivers that could establish such a bias on the lipid transfer.

## CONCLUSIONS

We developed a kinetic model for the fluorescence quenching experiments used to measure the rate of lipid transfer between vesicles mediated by tethering proteins. In these experiments, transfer of lipids labeled with fluorescence donors and acceptors to vesicles without labeled lipids leads to an overall dilution of labeled lipids. This dilution reduces the overall FRET efficiency, and therefore results in an increase in the fluorescence intensity at the donor emission frequency. In our theoretical model, lipid transfer occurs with a rate *k* through a single bottleneck formed by an Atg2 bridging between the vesicles. For dilute labeled lipids, the fluorescence signal then increases with a rate *r* = *k/N* where *N* is the total number of lipids in each of the outer leaflets. For *N* close to 11,000 for a vesicle of 50 nm diameter, the difference between the apparent rate *r* and the actual rate *k* is large.

For ATG2A-mediated lipid transfer, based on the experiments of Maeda et al. (2019), we obtained a lipid transfer rate of *k* ≈ 115 lipids per second and ATG2A. At this rate, ATG2A-mediated lipid transfer from the ER can contribute significantly to the growth of a phagophore on a minutes time scale. Already for a rather small bias of Δ*μ* = 1 *k*_B_*T*, thirty ATG2A molecules connecting the ER to the phagophore would supply one million lipids from the ER to the expanding phagophore within ten minutes. In our analysis of the data for yeast Atg2 reported by Osawa et al. (2019), the rate of lipid transfer appears to be even faster, about 750 lipids per second. On this basis, we conclude that Atg2-mediated lipid transfer from the ER to growing phagophores is kinetically feasible.

In addition to fast lipid transfer, we identified a second, slower phase in the lipid-transfer experiments by Maeda et al. (2019) and Osawa et al. (2019). Possible explanations include background signal, a sub-population of proteins with about 10-times slower transfer rates, lipid flip-flop, the formation of networks of Atg2-bridged vesicles or bleaching of fluorophores. A particularly appealing explanation is a subpopulation of Atg2 proteins in which the helix H4 has not been fully displaced from the lipid-transfer channel, as a likely prerequisite for fast lipid shuttling (Osawa et al. 2019) and as seen in a structure of Vps13 (Kumar et al. 2018). Irrespective of the physical origin of the slower phase, our quantitative kinetic analysis of the experiments shows that the bulk of the lipids is transferred rapidly, suggesting that the fast rate dominates also in a biological setting.

As a final point, we emphasize the fact that the kinetic model presented here is general. Beyond Atg2-mediated lipid transfer between labeled and unlabeled vesicles, it applies to a variety of experiments in which two reservoirs equilibrate with each other through a bottleneck. In particular, the model should prove useful for the quantitative analysis of measurements probing other proteins associated with lipid transfer at organelle contact sites, including the Mmm1-Mdm12 complex (Kawano et al. 2018) and VPS13 (Kumar et al. 2018). In addition, the model can be adapted to lipid shuttling between membranes via soluble lipid transfer proteins such as the Ups1-Mdm35 complex (Connerth et al. 2012, Watanabe et al. 2015). Whereas the interpretation of the rate *k* of lipid transfer changes, the calculation of the time-dependent FRET signal as a result of fluorescent lipid transfer remains unchanged.

## THEORY

### Atg2 shuttles lipids in single file

Atg2 presents a hydrophobic groove that was suggested to act as a conduit for phospholipids (Osawa et al. 2019). The hydrophobic groove along its long axis is so narrow that adjacent lipids are unlikely to exchange their position. In effect, this rules out regular diffusion of individual lipids along the groove as the transport mechanism. Instead, the structure of the N-terminal domain of Atg2 (Osawa et al. 2019) implies that single-file transport dominates lipid shuttling. As shown before for carbon nanotubes (Hummer et al. 2001) and formulated in a theory of single-file transport (Berezhkovskii and Hummer 2002), what diffuses back and forth is the boundary between particles (here: lipids) taken up from one or the other reservoir (here: the labeled and initially unlabeled vesicles). Based on their fluorescence measurements, Valverde et al. (2019) estimated that human ATG2A binds about *M* ≈ 20 lipids, which is roughly consistent with the 20-nm length of Atg2 (Osawa et al. 2019) and a spacing of 1 nm between the lipids. In the simplest model, the conduit thus holds *M* lipids in a single file. The boundary between the lipids from one and the other vesicle moves back and forth with a characteristic “hopping” time *τ*_hop_. We denote the rate of such hops “left” or “right” by one site, irrespective of direction, as *k*_hop_ = 1*/τ*_hop_. In every such hop, a lipid is pushed out of the conduit at one end and another lipid is sucked in at the other end. Whenever a lipid enters newly from one vesicle, it has a splitting probability of 1/(*M* + 1) to fully traverse the conduit and exit on the other side (Berezhkovskii and Hummer 2002). The effective rates of lipid transfer through such a single-file conduit is thus *k* = 1/[(*M* + 1)*τ*_hop_].

More elaborate models of single-file transport include the presence of vacant sites (Chou 1999). Indeed, for long single-file chains, we expect the dynamics of vacant sites to dominate the transfer process. However, single files with vacant sites will not alter the fundamental issue that all lipids pass through a single bottleneck (here: Atg2). Only the interpretation of the lipid transfer rate *k* in terms of hopping events and the scaling with the number of sites *M* will have to be adjusted.

### The population of labeled lipids relaxes with a rate that scales inversely with the total number of lipids

In the following, we develop a detailed kinetic model of lipid shuttling between two vesicles connected by a single Atg2 molecule. For now, we assume that lipid exchange occurs between the outer leaflets of the two vesicles. An extension to include lipid flip-flop between the leaflets is described below. We further assume that the two lipid vesicles have equal size. A generalization to vesicles differing in size will be given below. In addition, we assume that Atg2 transfers all lipids irrespective of type. In other words, the hopping rate of the chain is independent of which particular lipids it contains. In a biological setting, the latter assumption will have to be relaxed, at the very least by excluding certain lipids from transfer. However, here we concentrate on explaining the in vitro experiments with lipids containing similar tails. Finally, we assume that vesicle tethering by Atg2 is quasi-irreversible on the time scale of the experiments (i.e., two vesicles are either tethered for the duration of the experiment or not at all).

We denote the total number of lipids of type *j* (*j* = 1*, …, L* with *L* the number of distinct lipid types) in the outer leaflets of the two vesicles combined as *N_j_*, with *n_j_* lipids on vesicle 1 and *N_j_ − n_j_* lipids on vesicle 2. We denote the state of the two-vesicle system by the vector 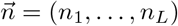. Let 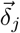 be a vector of zeros, with a one at position *j*. Transport of a lipid of type *j* from vesicle 2 to vesicle 1 then changes the state from 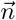 to 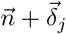, and the reverse process changes the state to 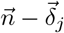. These lipid transport events are the elementary events in a kinetic master equation,

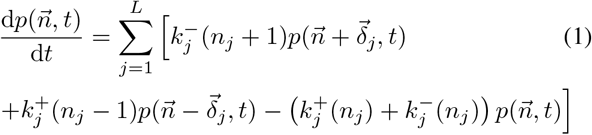

where 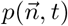 is the population of microstate 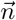 at time *t*, and 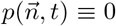 if any *n_j_ <* 0 or *n_j_ > N_j_* by definition. The rate coefficients 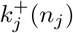 and 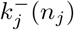 to increase and decrease the number of lipids of type *j* on vesicle 1 by one, respectively, are given by

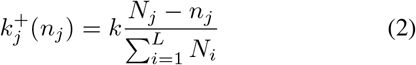

and

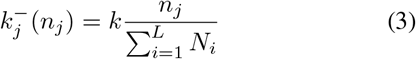

respectively. These rate coefficients satisfy a mass action law for vesicles of equal size and the condition that that the rate of transport events irrespective of direction and lipid type is exactly *k*,

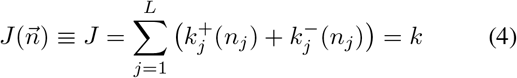

where the flux *J* = *k* going out of any microstate 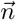 is independent of 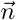. The single-file nature of the Atg2-mediated lipid transport enters the model through this condition on the rate. On average, we expect a single hopping event of the single-file chain to occur per hopping time, and we expect the chain hopping times to be exponentially distributed (Berezhkovskii and Hummer 2002).

At first sight, the master equation, Equation 1, with an exponential number of states,) 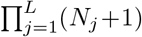, appears quite challenging. However, on closer inspection we realize that the probability can be factorized because each 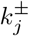 depends only on *n_j_* and not on the other elements of 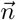. In other words, the dynamics of the lipid sub-populations are effectively independent of each other. We thus arrive at a factorized population

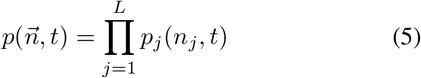

with *p_j_*(*n_j_*) = 0 if *n_j_ <* 0 or *n_j_ > N_j_*. The time-dependent *p_j_* satisfy a set of *L* one-dimensional master equations

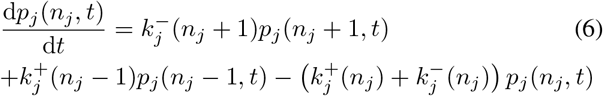

With the rate coefficients given in Equation 2 and Equation 3, Equation 6 is analytically tractable. The equilibrium populations are given by binomial distributions,

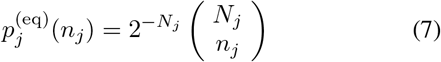

The eigenvalues of the rate matrix corresponding to Equation 6 are *λ*_0_ = 0*, λ*_1_ = −*r*, 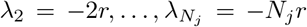 with

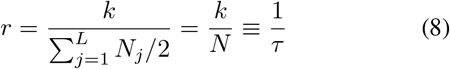

the lipid population relaxation rate. With equally spaced eigenvalues, Equation 6 corresponds to a kinetic harmonic oscillator. Here 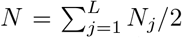 is the total number of lipids in an outer leaflet at equilibrium. *r* is the largest non-zero eigenvalue of the master equation and thus gives the rate of decay of the lipid populations by Atg2-mediated transport between the two vesicles. Indeed, by summing up the respective terms in Equation 6, one finds that the mean lipid number 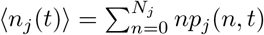 satisfies a simple rate equation and relaxes exponentially as

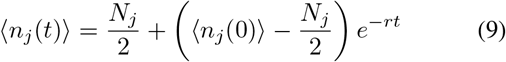

The variance in the number of lipids of type *j* grows as

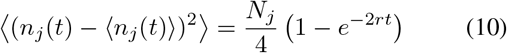

The expression for the mean number of lipids, Equation 9, corresponds exactly to the one obtained for the phenomenological rate model, Equation 32, with ⟨*n_j_*(0)⟩ = *N_j_* for the labeled vesicle. This model was constructed according to Figure 1. Importantly, the rate *r* and the characteristic time *τ* = 1*/r* of lipid population relaxation are independent of the lipid type and composition. However the relaxation rate is inversely proportional to *N*, i.e., the total number of lipids in a leaflet.

For completeness, we give expression for the left eigenvector of the rate matrix corresponding to eigenvalue *λ*_1_ = *−r*:

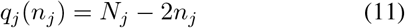

The vector of lipid counts *n_j_* is thus a linear combination of the left eigenvectors (1, …, 1) and *q_j_*(*n_j_*) for eigenvalues 0 and −*r*, respectively. Therefore, only the rate *r* appears in the relaxation of the mean population, Equation 9.

Finally, we point out that neither the full master equation, Equation 1, nor the corresponding one-dimensional master equations, Equation 6, put restraints on the total number of lipids in the outer leaflet of a vesicle. Instead, this number is itself binomially distributed with a mean of *N* and a variance of *N/*2. For vesicles larger than 20 nm in diameter, the relative fluctuations in the lipid number are less than 1% and thus negligible.

### Lipid transfer between vesicles of different size

The kinetic model can be readily extended to the case where the Atg2-tethered vesicles differ in size. We denote the total number of lipids on vesicles 1 and 2 with *M*_1_ and *M*_2_, respectively. At a particular instance in time, there are *n_j_* lipids of type *j* on vesicle 1 and *N_j_ −n_j_* on vesicle 2 with 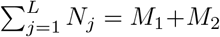. The rate coefficients 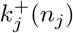 and 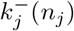 to increase and decrease *n_j_* by one are, respectively,

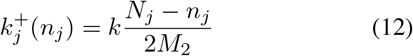

and

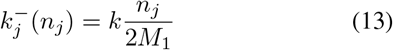

With these rate coefficients, the flux 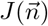 out of microstate 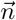 satisfies Equation 4, i.e., the net rate out of any microstate is 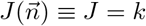 irrespective of state, as required for passage of all lipids through a single bottleneck. Again, we can factorize the corresponding master equation. The equilibrium population of the number of lipids of type *j* on vesicle 1 is again a binomial distribution,

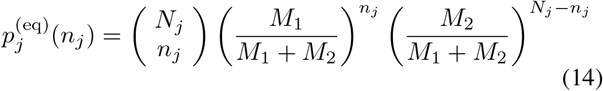

with mean *n_j_* = *N_j_M*_1_/(*M*_1_ + *M*_2_). The eigenvalues of the rate matrix corresponding to Equation 12 and Equation 13 are *λ*_0_ = 0*, λ*_1_ = *−r*, 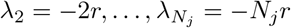 with

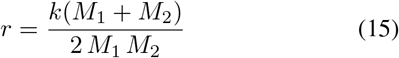

For equal-size vesicles with *M*_1_ = *M*_2_ = *N*, we recover the rate *r* = *k/N* of Equation 8.

### Dependence of the donor fluorescence on the number of donor and acceptor lipids on a vesicle

In the following, we calculate the expected change in the fluorescence signal resulting from transfer of labeled lipids between the two vesicles connected by an Atg2 conduit. Here we assume that the vesicles are (1) spherical and (2) small such that mixing of transferred lipids is fast compared to Atg2-mediated lipid transfer; that (3) FRET donor and acceptor lipids are dilute; and (4) that Förster theory applies. If a donor lipid is excited, the probability that it emits at the donor emission frequency is then one minus the probability of FRET to an acceptor lipid. The probability of FRET from a donor to an acceptor lipid *i* at Euclidian distance Δ_*i*_ is

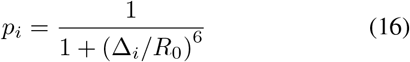

where *R*_0_ is the Förster radius. For simplicity, we assume here an ideal orientation factor of *κ*^2^ = 2/3. If we have *n* acceptor lipids in the vicinity, at distances Δ_*i*_ (*i* = 1*, …, n*), then the probability of emission at the donor frequency is given by the probability that no FRET occurs,

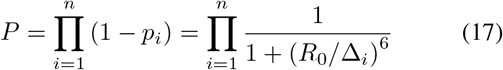

Here we treated the FRET events as independent, based on the assumption of dilute labeled lipids.

We now consider the case of a single donor lipid on a vesicle of radius *R*, together with *n* acceptor lipids that are uniformly distributed on the vesicle. The distribution of the Euclidian pair distances Δ is then *p*(Δ) dΔ = Δ dΔ/(2*R*^2^) for 0 ≤ Δ ≤ 2*R* and zero otherwise. If the donor lipid is excited, the probability to emit at the donor frequency is

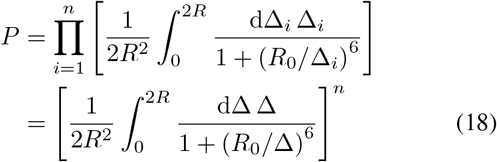

The integral can be evaluated analytically, giving us the quenching coefficient *Q* with *P* = (1 *− Q*)^*n*^ as

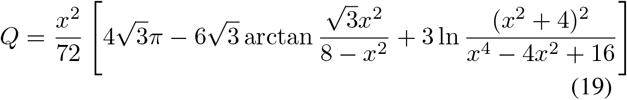

where *x* = *R*_0_*/R* is the ratio of Förster radius and vesicle radius. arctan denotes the inverse tangent. The range of the inverse tangent function for 0 *< x <* ∞ is 0 to 2*π/*3, as ensured in code by using the arctan2 function available in many programming languages. For small *R*_0_*/R*, we can approximate the quenching factor as

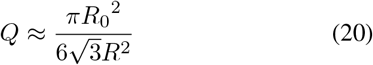

In realistic situations, we have *n* ≫ 1 and *R*_0_ ≪ *R*. Then we can replace (1 − *Q*)^*n*^ ≈ exp(−*nQ*). In this way, we arrive at the probability that a single donor fluorophore on a vesicle of radius *R* together is not quenched by *n* uniformly distributed acceptor fluorophores as

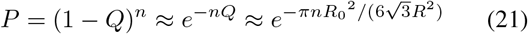

This is the probability that a single donor fluorophore emits at the donor frequency. The total fluorescence signal at the donor frequency with *m* donors is then proportional to *m* times *P*.

### The FRET fluorescence signal depends non-exponentially on time

We can now combine the results of the preceding two sections into a theoretical prediction of the overall fluorescence signal at the donor frequency as a function of time. In our simplest model of the experiments of Maeda et al. (2019), labeled vesicles containing 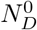 donor lipids, 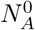 acceptor lipids and 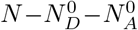 other lipids in their outer leaflets are mixed with equal-size unlabeled vesicles containing *N* unlabeled lipids each in their outer leaflets. The radius of these vesicles is *R*. Then, according to the master equation, Equation 1, we expect the populations of donor and acceptor lipids in the outer leaflets of the labeled (*L*) and unlabeled (*U*) vesicles to relax as

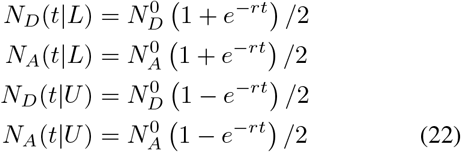

Here we assumed that lipid transfer is restricted to the outer leaflets. Generalizations to other types of initial conditions are straightforward.

By combining these time-dependent mean lipid numbers with the expressions for the fluorescence intensity given a certain number of lipids we arrive at an expression for the fluorescence intensity at the donor frequency as a function of time, summed over the labeled and initially unlabeled vesicles,

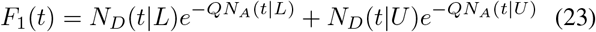

We also define a reduced signal in which we subtract the fluorescence at time zero (i.e., before mixing) and scale by the signal in the limit of long times,

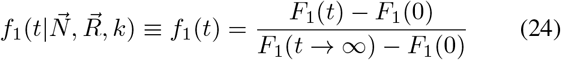

where 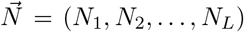 is the vector of lipid populations, 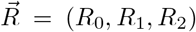 is the vector of Förster and vesicle radii (here: *R*_1_ = *R*_2_ = *R*), and *k* is the lipid transfer rate. The subscript 1 in *F*_1_(*t*) and *f*_1_(*t*) indicates that we account only for the outer leaflet, thereby neglecting the time-dependent effects of FRET between inner and outer leaflets. Below, we will extend the formulation to account also for the inner leaflets. The reduced signal *f*_1_(*t*) starts at zero and approaches one as *t* → ∞. However, it is not a simple exponential, containing terms of the form exp(−*α −rt*+*γ* exp(−*rt*)). As a consequence, the rise in the signal is initially faster and then slows down compared to an exponential with the same characteristic time. For values of 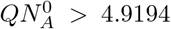, the reduced signal overshoots one at intermediate times and then drops back to one as *t* → ∞. A non-monotonic fluorescence signal in vesicle mixing experiments has indeed been reported by Kawano et al. (2018) in Figure 5C of their study of lipid transfer mediated by the Mmm1–Mdm12 complex.

### Interleaflet FRET

With the thickness of lipid bilayers and typical Föster radii being comparable in length, we expect a small contribution to the fluorescence quenching to come from interleaflet FRET. The distribution of pair distances Δ between points distributed randomly on two concentric spheres of radius *R*_1_ and *R*_2_, respectively, is *p*(Δ) dΔ = Δ dΔ/(2*R*_1_*R*_2_) for |*R*_1_ − *R*_2_| ≤ Δ ≤ *R*_1_ + *R*_2_ and zero otherwise. With this distribution, we can calculate the probability *P* = (1 − *Q*)^*n*^ for an acceptor dye on one of the two spheres not to be quenched by *n* donor dyes randomly distributed on the other sphere according to Equation 18. In this way, we obtain a quenching factor *Q* for FRET between leaflets,

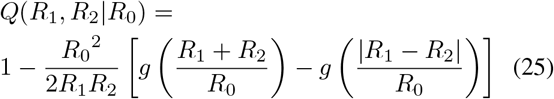

with

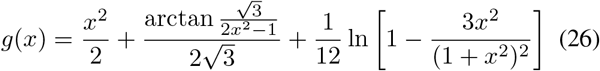

As in Equation 19, the range of the inverse tangent function for 0 *< x <* ∞ is 0 to 2*π/*3. In the limit of equal-size spheres, *R*_2_ = *R*_1_, we recover Equation 19.

In a fluorescence quenching experiment probing lipid transfer between the outer leaflets of a labeled and an initially unlabeled vesicle, we can now include the effect of lipids on the inner leaflets. Let 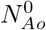 be 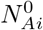 be the number of acceptor lipids in the outer and inner leaflets of the labeled vesicles, respectively, and 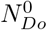 and 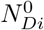 the corresponding number of donor lipids. For the time-dependent lipid numbers in the outer leaflets (*o*), we use Equation 22, with 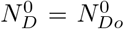 and 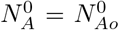. We assume the inner leaflets (*i*) not to mix and accordingly set

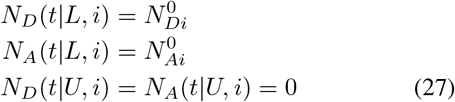

Adjusted to account for both intraleaflet and interleaflet FRET, Equation 23 for the expected fluorescence intensity becomes

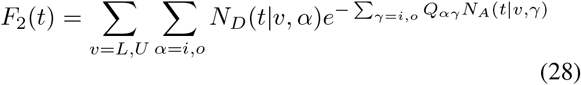

The sums are over the inner and outer leaflets, *i* a nd *o*, of labeled (*L*) and unlabeled vesicles (*U*). The quenching factors are *Q_oo_* = *Q*(*R*_1_*, R*_1_ | *R*_0_), *Q_oi_* = *Q_io_* = *Q*(*R*_1_*, R*_2_ | *R*_0_) and *Q_ii_* = *Q*(*R*_2_*, R*_2_ | *R*_0_). For lipid bilayers, we would roughly estimate *R*_1_ = *R*_2_ + *h* where the effective membrane thickness *h* defined by the typical dye positions is approximately 4 to 5 nm, depending on lipid composition, fluorescence dyes, and dye linkers. In our analysis, we used a value of *h* = 4.5 nm. Substituting the time-dependent lipid populations of Equation 22 and Equation 27 into Equation 28, we find

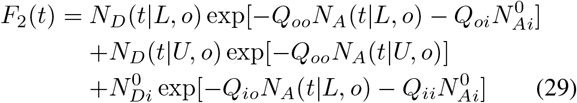

Depending on the ratio of the effective membrane thickness *h* = |*R*_1_ − *R*_2_| to the Förster radius *R*_0_, accounting for quenching by the acceptor lipids in the inner leaflets may be required. As in Equation 24, we define a scaled function

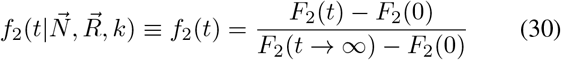

that starts at zero and approaches 1 as *t → ∞*. Note that the vector 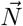 of lipid numbers defined above is expanded to include inner leaflets.

### Phenomenological rate models of lipid transfer

Bypassing the construction of a master equation for the transfer of individual lipids, we can instead invoke a macroscopic rate law. In this simplified model, we account for the fact that we are monitoring only labeled lipids by setting the apparent rate of lipid transfer as *r\** = *k/*(2*N*) = *r/*2 where 2*N* is the total number of lipids in the two outer leaflets. Here, scaling with the total number of lipids 2*N* accounts for the fact that only a fraction of the lipids is labeled and therefore tracked (see schematic in Figure 1). The factor of 1/2 accounts for the fact that only half of the transfers are in a given direction. The average number of lipids of type *j* in the outer leaflets of the initially labeled and unlabeled vesicles, *O*_1_(*t*) and *O*_2_(*t*), then evolve according to the first-order rate model

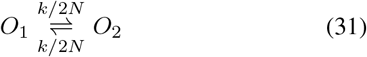

With 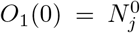 and 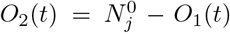 for *j* = *A* (acceptor lipids) or *D* (donor lipids), we arrive at

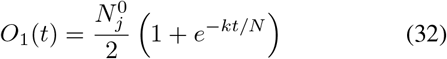

With this phenomenological rate model, we thus recover the solution for the mean number of lipids, Equation 9, of the full master equation.

To account for lipid flip-flop, we extend the rate model Equation 31 to include interleaflet lipid exchange,

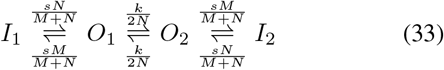

where *I_j_* and *O_j_* are the number of labeled lipids in the inner and outer leaflets of vesicles *j* = 1, 2, *s* is the flip-flop rate, and *N* and *M* are the total number of lipids in outer and inner leaflets, respectively. In the absence of lipid transfer between vesicles (i.e., *k* = 0), the populations of labeled lipids in inner and outer leaflets relax with rate *s* to a population ratio of *I*_1_*/O*_1_ = *I*_2_*/O*_2_ = *M/N*.

The analytical solutions of the first-order rate equations corresponding to Equation 33 with initial conditions *I*_1_(0) = *ρM*, *O*_1_(0) = *ρN* and *I*_2_(0) = *O*_2_(0) = 0 were obtained for donor and acceptor lipids, with *ρ* their respective fraction of the total lipid number on the labeled vesicle at time zero. The analytical expressions for the mean lipid numbers as function of time were combined with Equation 28 for the fluorescence intensity at the donor frequency to model the data in Figure S7.

## ACKNOWLEDGMENTS

The authors thank Sascha Martens and Jakob Bullerjahn for insightful discussions and acknowledge funding by the Human Frontier Science Program (RGP0026/2017) and the Max Planck Society.

---

**Supporting Figure S4.**
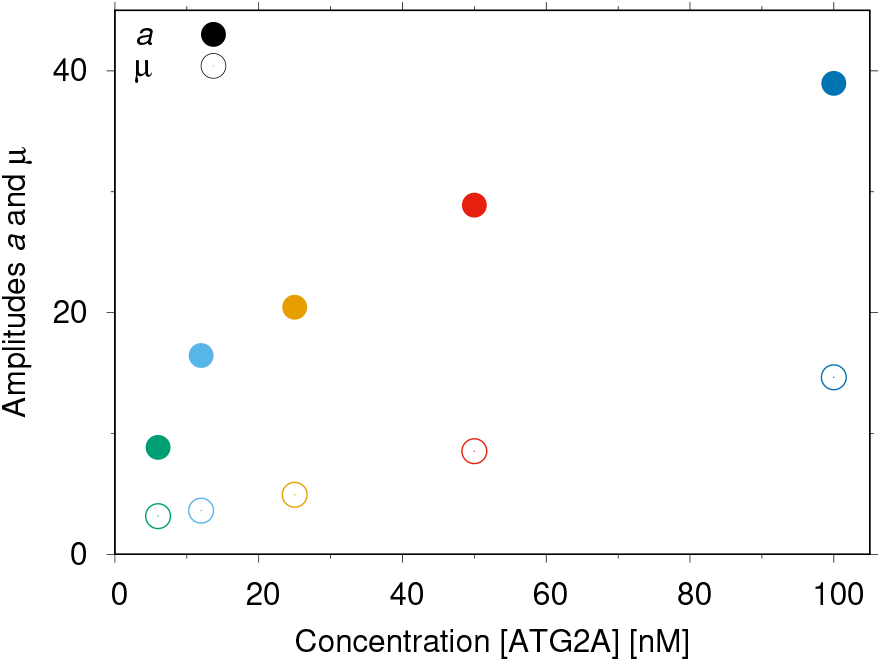
Amplitudes *a* and *μ* of signal and background, respectively, in fit of two-leaflet model 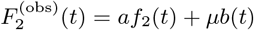 to fluorescence signal in Figure 3C of Maeda et al. (2019), as shown in Figure 2A.

**Supporting Figure S5.**
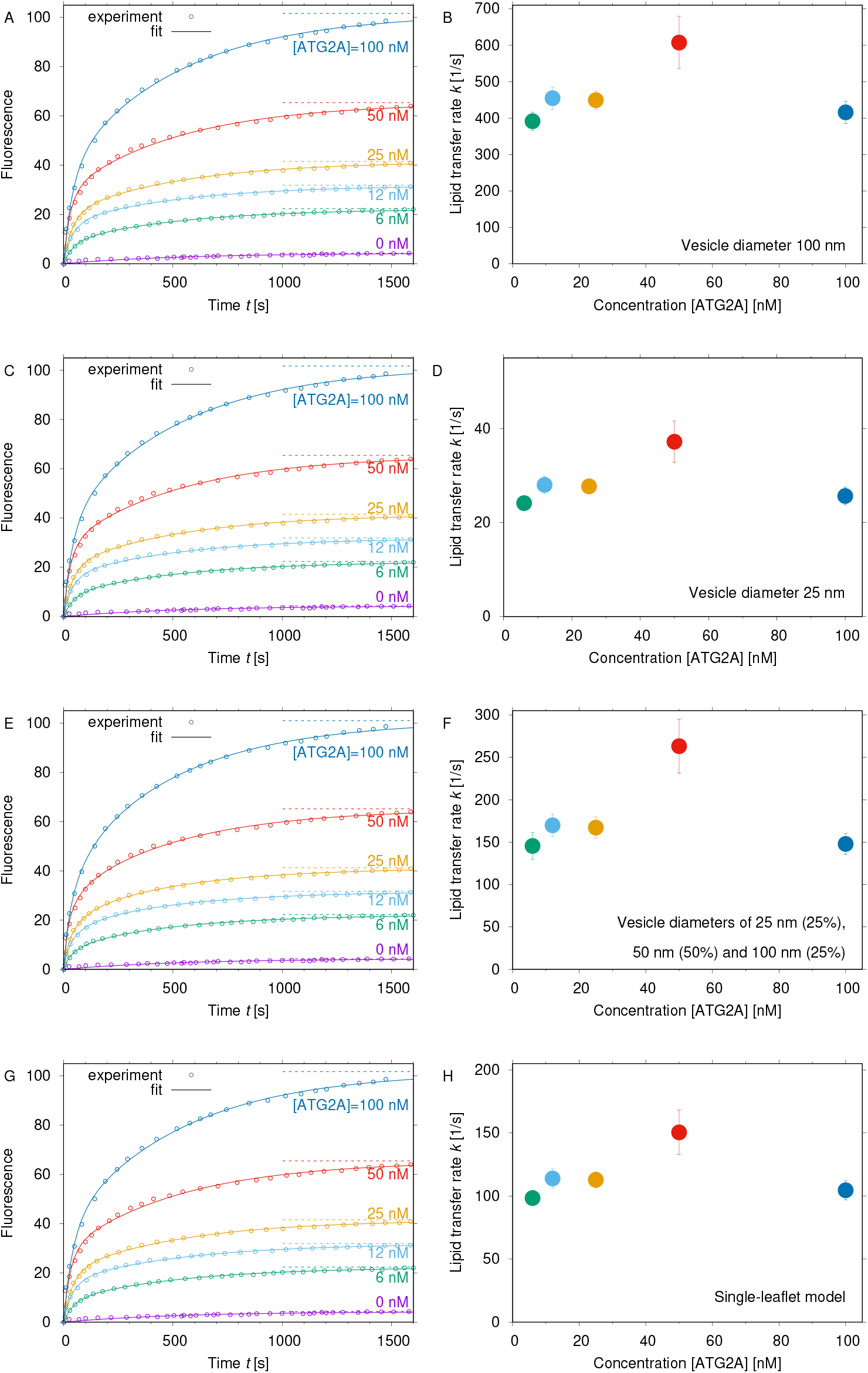
Rate of ATG2A-mediated lipid transfer from fits of kinetic model to FRET lipid-transfer measurements by Maeda et al. (2019) as in Figure 2. (A,B) Fit of two-leaflet model to fluorescence signal (A) and resulting lipid-transfer rate *k* (B) for vesicles with a diameter of 100 nm. (C,D) Fit of two-leaflet model to fluorescence signal (C) and resulting lipid-transfer rate *k* (D) for vesicles with a diameter of 25 nm. (E,F) Fit of two-leaflet model to fluorescence signal (E) and resulting lipid-transfer rate *k* (F) for a mixture of vesicles with diameters of 25 nm (25%), 50 nm (50%) and 100 nm (25%). (G,H) Fit of single-leaflet model to fluorescence signal (G) and resulting lipid-transfer rate *k* (H) for vesicles with a diameter of 50 nm. Horizontal dashed lines indicate the limiting values as time goes to infinity.

**Supporting Figure S6.**
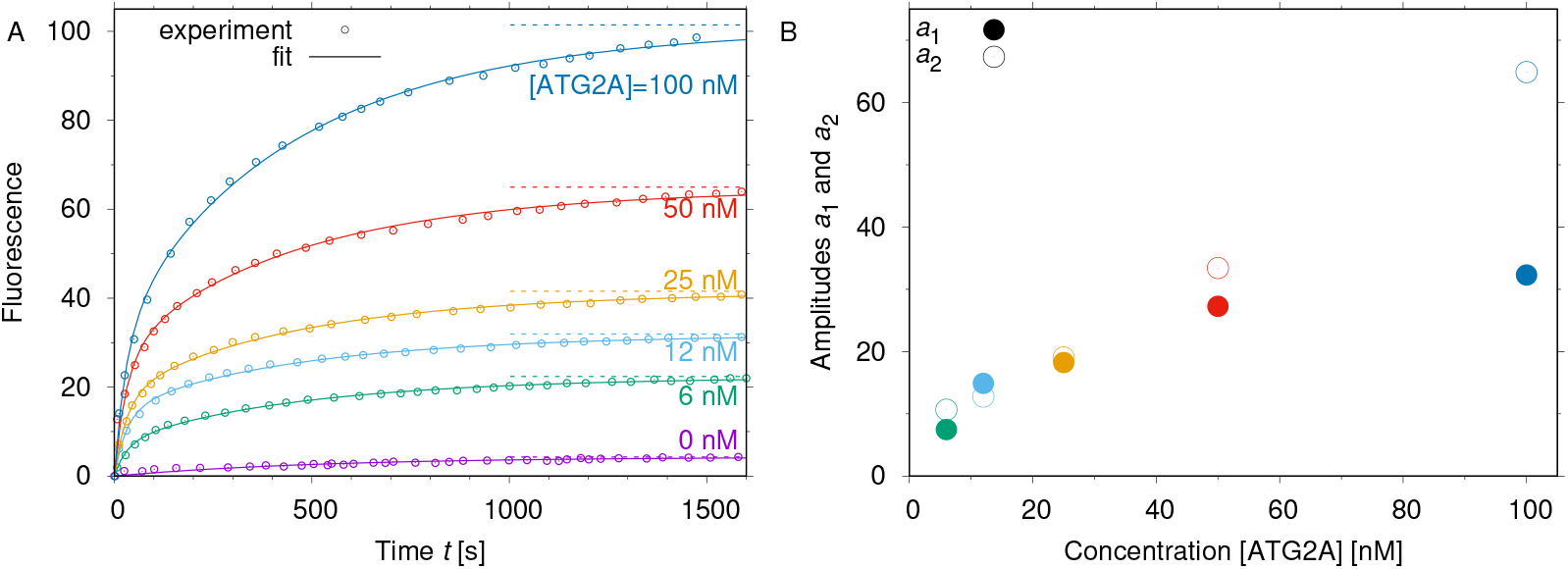
Global fit of two-population kinetic model 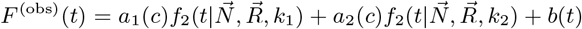 to FRET lipid-transfer measurements reported in Figure 3C of Maeda et al. (2019). Horizontal dashed lines indicate the limiting values as time goes to infinity. (A) Fits of the two-population model to the fluorescence traces. (B) Amplitudes *a*_1_ (filled symbols) and *a*_2_ (open symbols) of the fast and slow process with fitted rates *k*_1_ = 142 s^−1^ and *k*_2_ = 10 s^−1^, respectively. The fits are for a vesicle diameter of 50 nm.

**Supporting Figure S7.**
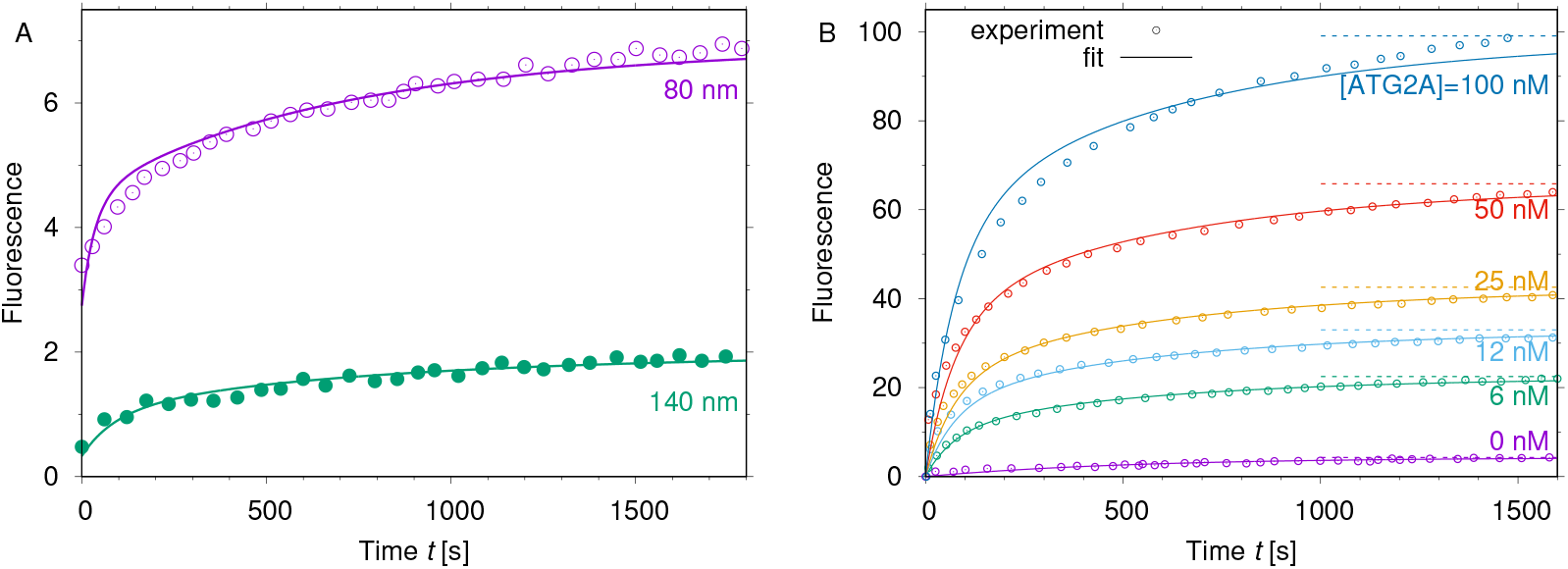
Global fit (lines) of lipid transfer kinetics with lipid flip-flop. (A) Results for yeast Atg2 using experimental data (symbols) taken from Figure 3f of Osawa et al. (2019) (80 nm diameter: magenta; 140 nm diameter: green). (B) Results for human ATG2A using experimental data (symbols) taken from Figure 3C of Maeda et al. (2019).

## References

Armstrong VT, Brzustowicz MR, Wassall SR, Jenski LJ, Stillwell W. Rapid flip-flop in polyunsaturated (docosahexaenoate) phospholipid membranes. Archives of Biochemistry and Biophysics. 2003; 414(1):74–82.

Berezhkovskii A, Hummer G. Single-file transport of water molecules through a carbon nanotube. Physical Review Letters. 2002; 89(6):064503.

Chou T. Kinetics and thermodynamics across single-file pores: Solute permeability and rectified osmosis. Journal of Chemical Physics. 1999; 110(1):606–615.

Chowdhury S, Otomo C, Leitner A, Ohashi K, Aebersold R, Lander GC, Otomo T. Insights into autophagosome biogenesis from structural and biochemical analyses of the ATG2A-WIPI4 Complex. Proceedings of the National Academy of Sciences USA. 2018; 115(42):E9792–E9801.

Connerth M, Tatsuta T, Haag M, Klecker T, Westermann B, Langer T. Intramitochondrial transport of phosphatidic acid in yeast by a lipid transfer protein. Science. 2012; 338(6108):815–818.

Hummer G, Rasaiah JC, Noworyta JP. Water conduction through the hydrophobic channel of a carbon nanotube. Nature. 2001; 414(6860):188–190.

Kalra A, Garde S, Hummer G. Osmotic water transport through carbon nanotube membranes. Proceedings of the National Academy of Sciences USA. 2003; 100:10175–10180.

Kawano S, Tamura Y, Kojima R, Bala S, Asai E, Michel AH, Kornmann B, Riezman I, Riezman H, Sakae Y, Okamoto Y, Endo T. Structure-function insights into direct lipid transfer between membranes by Mmm1-Mdm12 of ERMES. Journal of Cell Biology. 2018; 217(3):959–974.

Kumar N, Leonzino M, Hancock-Cerutti W, Horenkamp FA, Li PQ, Lees JA, Wheeler H, Reinisch KM, De Camilli P. VPS13A and VPS13C are lipid transport proteins differentially localized at ER contact sites. Journal of Cell Biology. 2018; 217(10):3625–3639.

Lantzsch G, Binder H, Heerklotz H. Surface area per molecule in lipid/C_12_E*n* membranes as seen by fluorescence resonance energy transfer. Journal of Fluorescence. 1994; 4(4):339–343.

Lee BC, Khelashvili G, Falzone M, Menon AK, Weinstein H, Accardi A. Gating mechanism of the extracellular entry to the lipid pathway in a TMEM16 scramblase. Nature Communications. 2018; 9:3251.

Maeda S, Otomo C, Otomo T. The autophagic membrane tether ATG2A transfers lipids between membranes. eLife. 2019; 8:e45777.

Malvezzi M, Andra KK, Pandey K, Lee BC, Falzone ME, Brown A, Iqbal R, Menon AK, Accardi A. Out-of-the-groove transport of lipids by TMEM16 and GPCR scramblases. Proceedings of the National Academy of Sciences USA. 2018; 115(30):E7033–E7042.

Osawa T, Ishii Y, Noda NN. Human ATG2B possesses a lipid transfer activity which is accelerated by negatively charged lipids and WIPI4. Genes to Cells. 2020; 25(1):65–70.

Osawa T, Kotani T, Kawaoka T, Hirata E, Suzuki K, Nakatogawa H, Ohsumi Y, Noda NN. Atg2 mediates direct lipid transfer between membranes for autophagosome formation. Nature Structural and Molecular Biology. 2019; 26(4):281–288.

Pan J, Tristram-Nagle S, Kucerka N, Nagle JF. Temperature dependence of structure, bending rigidity, and bilayer interactions of dioleoylphosphatidylcholine bilayers. Biophysical Journal. 2008; 94(1):117–124.

Siggel M, Bhaskara RM, Hummer G. Phospholipid scramblases remodel the shape of asymmetric membranes. Journal of Physical Chemistry Letters. 2019; 10(20):6351–6354.

Struck DK, Hoekstra D, Pagano RE. Use of resonance energy transfer to monitor membrane fusion. Biochemistry. 1981; 20(14):4093–4099.

Valverde DP, Yu SL, Boggavarapu V, Kumar N, Lees JA, Walz T, Reinisch KM, Melia TJ. ATG2 transports lipids to promote autophagosome biogenesis. Journal of Cell Biology. 2019; 218(6):1787–1798.

Watanabe Y, Tamura Y, Kawano S, Endo T. Structural and mechanistic insights into phospholipid transfer by Ups1-Mdm35 in mitochondria. Nature Communications. 2015; 6:7922.

